# An *in cell* site-specific labeling methodology reveals conformational changes of proteins in bacteria

**DOI:** 10.1101/2022.08.03.502720

**Authors:** Yulia Shenberger, Lada Gevorkyan-Airapetov, Melanie Hirsch, Lukas Hofmann, Sharon Ruthstein

## Abstract

Gaining new structural information on proteins in their native cellular environments will shed light on many enzymatic reaction mechanisms and encourage the development of new therapeutic approaches. During the last decade, *in cell* electron paramagnetic resonance (EPR) spectroscopy experiments have provided high-resolution data on conformational changes of proteins within the cell. However, one of the major obstacles of EPR spectroscopy is the spin-labeling process, which until now was performed only outside the cellular environment (i.e., exogenously). The spin-labeled protein is then injected into the cell, which limits the protein size and the cellular system that can be used. Here, we describe a new spin-labeling approach that can be applied to over-expressed proteins in *Escherichia coli* (i.e., endogenously). This approach uses a Cu(II) ion bound to a ligand, which has high affinity to a dHis site in the protein of interest. The presence of a nearby ^19^F-phenylalanine residue can be exploited to verify that the Cu(II)-ligand indeed bound to the protein target. This new methodology allows for the study of any protein, regardless of size or the cellular system used.

---

Understanding structural dynamics of proteins within their physiological environment can be considered as the Holy Grail of structural biology. Currently, the most common methods used for determining protein structure in the cell are nuclear magnetic resonance (NMR), Forster resonance energy transfer (FRET), and cryo-electron microscopy (cryo-EM). These fast-advancing technologies provide different perspectives on the spatial-temporal resolution and structural rearrangements of proteins within the cell. *In cell* NMR is capable of detecting interactions between proteins and small molecules ^*1, 2*^, yet is limited by the size of the biological system of interest. *In cell* FRET can report on dynamical changes of proteins in the cell ^*3, 4*^ but cannot provide accurate information on topological changes that occur upon structural rearrangements. *In cell* cryo-EM is an excellent tool for obtaining structural information on large symmetric biological systems and complexes. At the same time, it is less preferred for monitoring proteins of low abundance or low symmetry ^*5*^. In the last decade, *in cell* electron paramagnetic resonance (EPR) spectroscopy has emerged as an excellent methodology to follow biological mechanisms within the cell ^*6–12*^. EPR distance measurements, such as double electron electron resonance (DEER), can define distances within a biological system in the nanometer range (1.5-10.0 nm) ^*13*^.

**Scheme 1.**
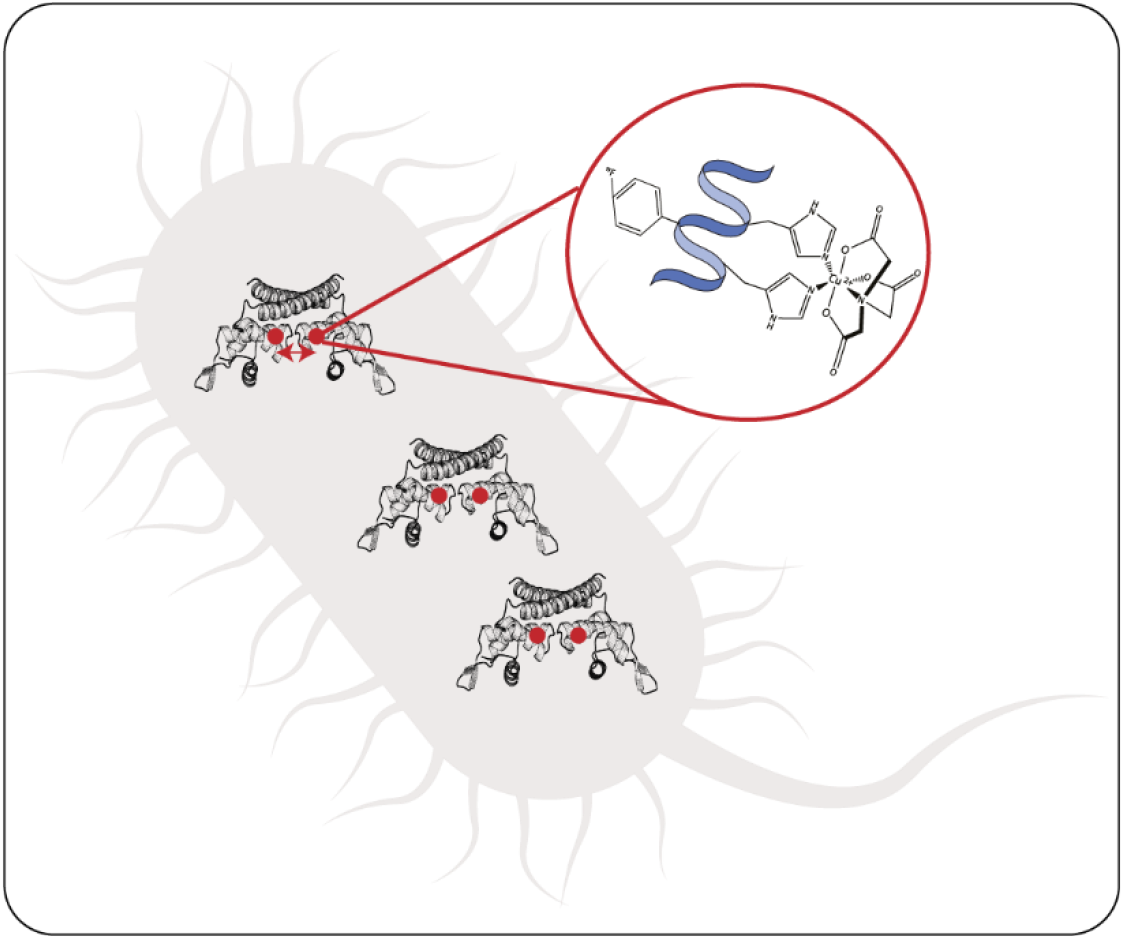
Schematic depiction of the spin-labelling approach. The protein chosen here is the CueR_^19^F58_L60H_G64H mutant (PDB 1Q05) over-expressed in *E. coli*. The protein is characterized by a dHis site (residues L60H and G64H) and ^19^F-labeled phenylalanine (residue F58). Cu(II) coordinated to a nitriloacetic acid (NTA) ligand, which displays high affinity to the dHis site, is injected into the cells to realize site-specific labelling within the cell.

EPR spectroscopy offers numerous advantages, relative to other biophysical tools. First, is its high sensitivity, EPR can target biomolecules present at concentrations ranging from the micromolar level to the few tens nanomolar level. Moreover, EPR measurements are not limited by the size or complexity of the biological system nor by the environment in which it is found. However, EPR spectroscopy requires paramagnetic centers, a need that raises several challenges, especially for *in cell* EPR experiments. The first obstacle is that the selected spin-label should be stable in a reduced environment, such as the cytoplasm. Therefore, the most common spin-labels used for *in cell* EPR measurements are Gd(III) ^*14*^- and trityl-^*15–18*^ based spin-labels. A second obstacle is that the spin-labeling methodology currently in use requires injection of the spin-labeled protein into the cell, after the spin-labeling procedure was performed outside the cell. This limits the size of the biomolecule that can be studied, as well as the cellular system that can be used, which is limited by how much the cell membrane can be distorted. As such, this method is mostly employed for studies of eukaryotic cellular systems.

We now report the development of a new *in cell* spin-labeling methodology. The advantage of this method is that it is performed on over-expressed proteins within the bacterial cell, with the spin-labeling process being carried out within the cell itself. Using this method, any protein, regardless of size and/or complexity, can be studied in a variety of cellular systems. In this approach, we use Cu(II)-nitriloacetic acid (Cu(II)-NTA) as the spin-label. Cu(II)-NTA shows high-affinity to dHis sites, especially for dHis sites located within helices (Scheme 1) ^*19–22*^. Moreover, the NTA ligand ensures that Cu(II) will not be reduced in the reducing environment of the cell. The two histidine residues of such sites should be separated by four amino acids to ensure high Cu(II)-NTA binding. Since this spin-labeling method is performed on the protein backbone, it provides highly narrow distance distribution functions, and can differentiate between minor conformational changes of the protein. However, to ensure that Cu(II)-NTA molecules reach the specific dHis site of the over-expressed protein and do not bind to other His-based sites that may exist in the protein, a ^19^F-phenylalanine close to the dHis site is introduced. ^19^F-phenylalanine can serve to quantify the ratio between the Cu(II)-NTA spin-labeling and the targeted protein.

As proof of concept of this methodology, the *Escherichia coli (E. coli)* copper-sensitive transcription factor CueR ^*23–29*^ was employed. CueR is a metalloregulator protein that prevents copper toxicity in Gram-negative bacteria. It possesses high affinity to the reduced form of copper, Cu(I). Upon copper binding, CueR initiates a transcription process that leads to the appearance of proteins that either oxidize copper to the less toxic Cu(II) form or shuttle copper outside the cell. We previously showed that DEER measurements performed on a CueR_L60H_G64H mutant labeled with Cu(II)-NTA can follow conformational changes of the protein as a function of Cu(I) and DNA binding ^*24*^. Next to the L60H_G64H, a phenylalanine residue exists (F58). So as to not to interfere with the secondary structure of the protein, we replaced this residue with ^19^F-phenylalanine. Therefore, for *in cell* EPR measurements, the CueR_^19^F58_L60H_G64H mutant was over-expressed in *E. coli*. The expression and purification protocols are described in the SI. SDS-PAGE confirmed CueR over-expression in the cell, as well as the purity of purified CueR_^19^F58_L60H_G64H (Fig. S1, SI). Circular dichroism showed that the secondary structure of the protein was not affected by this mutation (Fig. S2, SI). An electrophoresis mobility shift assay (EMSA) confirmed that the CueR_^19^F58_L60H_G64H mutant protein bound to the *copA* promoter in a similar manner as the native CueR protein (Fig. S3, SI). Assessing cell viability in the presence of Cu(II)-NTA and free Cu(II) ions at various concentrations (Fig. S4, SI) revealed that at the concertation used in this study, cell viability was not affected. To ensure that free Cu(II) did not provide any EPR signal, continuous wave EPR experiments were carried out at 130K with 50 μM and 100 μM CuCl_2_ being inserted into the *E. coli* cells. The EPR spectra (Fig. S5, SI) did not show any signal, confirming that all free Cu(II) ions were reduced.

Hyperfine pulsed EPR methods that evaluate the interaction between paramagnetic centers and nearby nuclei were applied to quantify the amount of Cu(II)-NTA near ^19^F nuclei. Since the distance between Cu(II) and ^19^F is expected to be in the 10-15 Å range, and given the comparable short relaxation time (Fig. S6, SI), electron nuclear double resonance (ENDOR) experiments could not detect this interaction. However, electron spin echo envelope modulation (ESEEM) experiments observed the 42 MHz peak in the Fourier transform (FT) spectrum of ESEEM performed in Q-band frequency (Fig. S7, SI). Q-band ESEEM experiments also succeeded in differentiating between the ^19^F and ^1^H peaks in the FT spectrum (Fig. 1a). Repeating the experiment in deuterated buffer led to a reduced ^1^H peak. Moreover, the FT spectrum showed close binding to His residues, exhibiting characteristic peaks of ^14^N nuclei in the ESEEM FT spectrum (see Fig. S7, SI) ^*30–33*^.

**Figure 1.**
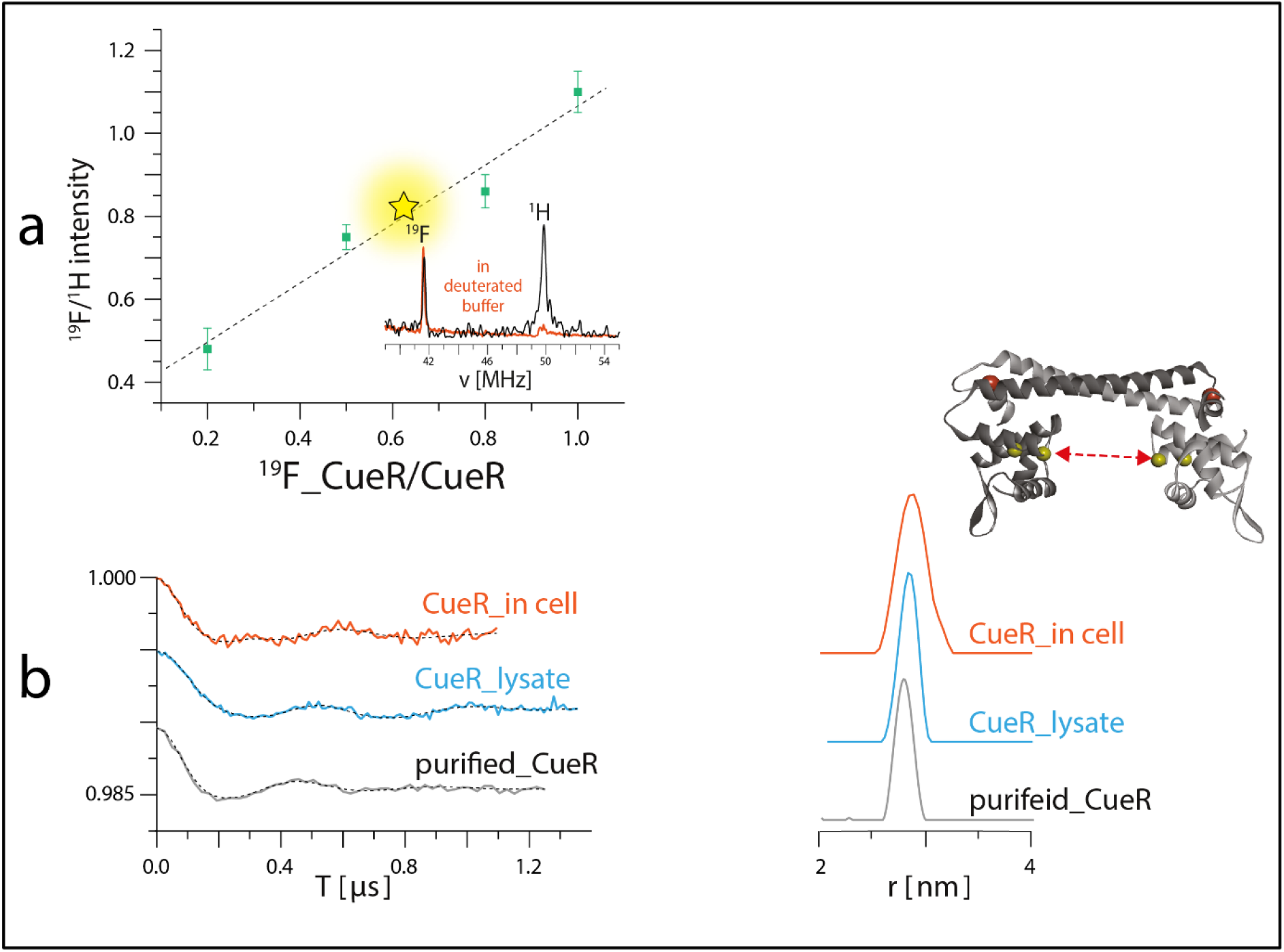
EPR measurements on *E. coli* CueR_^19^F58_L60H_G64H spin-labelled with Cu(II)-NTA. **a**. The intensity of ^19^F vs ^1^H from the Q-band 3P-ESEEM FT for different ratios of CueR_L60H_G64H labelled with ^19^F58 (CueR_^19^F58_ L60H_G64H) to non-labelled CueR_F58_ L60H_G64H. The yellow star corresponds to the ratio detected in the cell. In deuterated buffer the ^1^H peak is educed (inset red line). **b**. Q-band EPR distance measurements on purified (gray line) and overexpressed protein lysate (light blue) and in the cell (orange). CueR is a dimer, and the distance is measured between the two Cu(II)-dHis sites (inset).

To quantify the amount of Cu(II)-NTA next to ^19^F nuclei, we mixed CueR_^19^F58_L60H_G64H with non-isotope labeled phenylalanine CueR_L60H_G64H at different ratios. Figure 1a shows the changes in ^19^F/^1^H intensity in Q-band ESEEM FT spectra at different ratios of ^19^F-labeled and non-labeled *E. coli* CueR_L60H_G64H proteins. The linear correlation observed confirmed that this methodology can be used to quantify the amount of Cu(II) site-specifically bound to a protein. We then relied on ESEEM experiment to evaluate the amount of Cu(II) bound to CueR_^19^F58_L60H_G64H over-expressed in *E. coli*. For different independent experiments, we found that the ratio between ^19^H/^1^H suggested 60-70% binding of the Cu(II) to the protein. This value is consistent with the change in modulation depth values between purified CueR (0.006 ± 0.001) and CueR detected in the cell (0.0045 ± 0.001). This ratio allows a minimal background contribution (see Fig. S8, SI) to the DEER signal, as reported by Ketter et al. ^*34*^. Furthermore, it confirms that for this biological system, there is no additional high-affinity site to which Cu(II)-NTA can bind.

DEER experiments were run on three samples, namely, CueR_^19^F58_L60H_G64H over-expressed in *E. coli* cells, *E. coli* lysate and purified CueR_^19^F58_L60H_G64H. Cu(II)-NTA was added to the cells together with the isopropyl-β-D-thiogalactopyranoside inducer (first sample). To test for promotor leakiness, the cells were harvested, and the cleared growth medium was measured by EPR. This revealed no Cu(II) signal (Fig. S5, SI). Subsequently, the harvested cells were lysed, the cell debris was collected by centrifugation, and clear lysate was measured (second sample). Finally, the data collected were compared to that obtained with the purified protein (third sample). The field-sweep echo-detected EPR spectra for all three samples are shown in Fig. S6, SI. For *in cell* and lysate experiments, a Mn(II) signal was noted. However, the position of pulses chosen were in the g_⊥_ region, which is far from Mn(II) in the Q-band. Altogether, the distance distribution for the purified protein was 2.7 ± 0.1 nm, whereas in the lysate it was 2.8 ± 0.1 nm yet was slightly broader (2.85 ± 0.25 nm) in the cell. The validation of the distance distribution function is reported in Fig. S9, SI. The agreement between the data confirms that most of the Cu(II)-NTA successfully bound to the over-expressed protein within the cell.

*In cell* conformational changes of CueR were monitored as a function of Cu(I) binding. Here, DEER measurements were employed in presence of free copper. As noted above, when cells are incubated with CuCl_2_, Cu(II) is immediately reduced to Cu(I) in the cellular environment (Fig. S5, SI). We added 50 μM of CuCl_2_ to cells over-expressing CueR_^19^F58_L60H_G64H, with such over-expression being induced in presence of Cu(II)-NTA. Comparison of the DEER data collected *in cell* with that for purified holo_CueR at a ratio of 1:1 Cu(I):CueR (Fig. 2) showed a minor shift in the average distance seen for the purified protein, collected *in vitro*, and for the protein assessed *in cell* (2.4 ± 0.2 nm and 2.2 ± 0.2 nm, respectively). Since it is difficult to quantify CueR-bound Cu(I), as compared to the data collected from the protein *in vitro*, this difference might be attributed to different Cu(I) ions that coordinate CueR *in vitro* and in the cell. We recently suggested that each CueR monomer can coordinate two Cu(I) ions, and that an excess of copper induces different conformational and dynamical changes in CueR ^*35*^. Previously, molecular dynamics simulations carried out on CueR suggested two modes of motions, specifically, bending and twisting upon Cu(I) binding ^*36*^. The EPR data pointed to the two α4 helices getting closer to each other upon Cu(I) binding (Fig. 3). Such motion would reduce the distance between the two DNA-binding domains, thereby inducing transcription initiation, as proposed previously ^*24, 25, 35*^.

**Figure 2.**
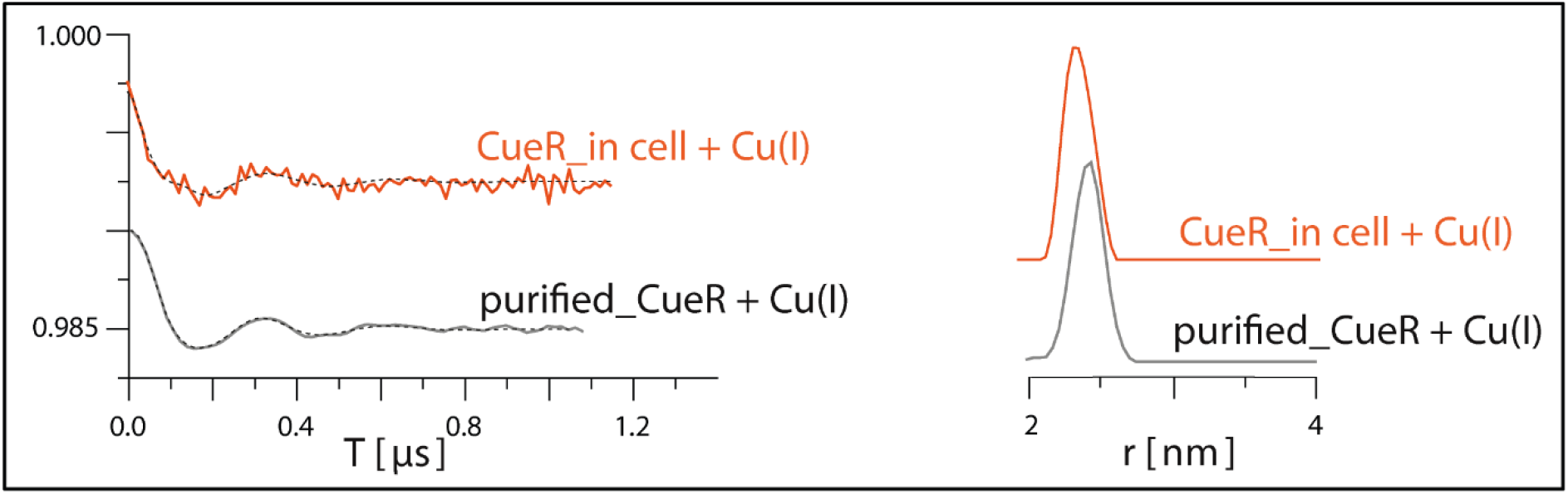
EPR measurements of *E. coli* CueR_^19^F58*_*L60H_G64H spin-labelled with Cu(II)-NTA in the presence of Cu(I). Q-band EPR distance measurements of purified (gray line) and over-expressed protein in the cell (orange) in the presence of Cu(I) ions.

**Figure 3.**
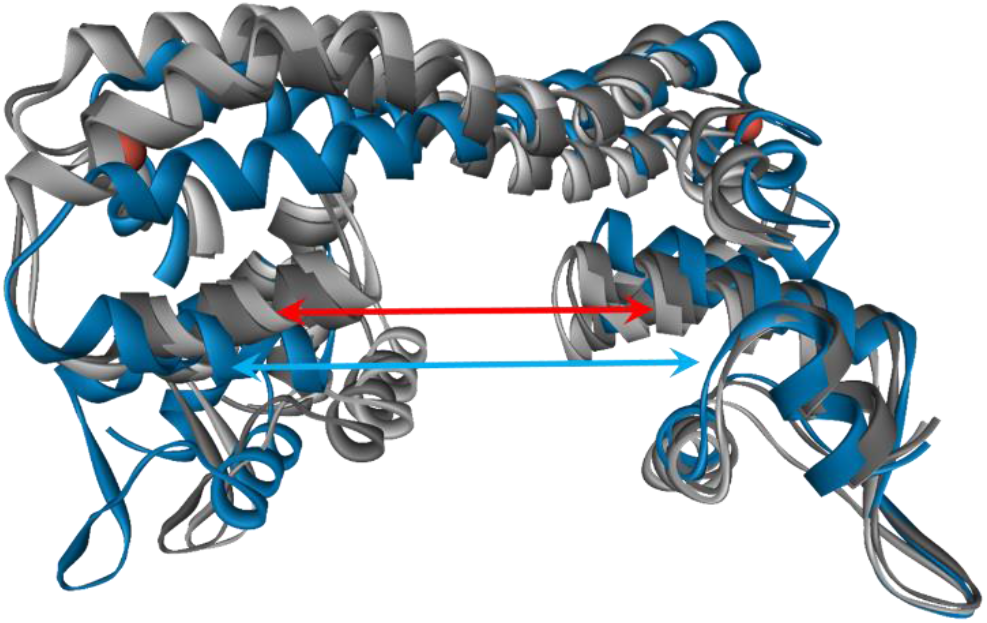
Structural models of CueR. CueR in the apo-form (PDB 1Q05, blue color), and in the two active structures (grey colors) proposed to exist in the presence of copper and reported in Sameach at al.^*24*^ are shown. Changes in distance between the two α4 helices are marked by arrows (blue arrow for the apo state, red arrow for the holo state).

Here, we showed the feasibility of using an endogenously over-expressed protein in *E. coli* and characterized by a dHis site to spin-label the protein with a Cu(II) paramagnetic center in the cellular environment. The main advantage of this method is that it can be performed on any over-expressed bacterial protein, independent of its size or complexity. The labeling yield is comparable high at about 60%, which allows for the detection of well-resolved DEER time-domain signals. It is important to note that the experiments were performed at low temperature (i.e., 20K), and required long acquisition times of several days in order to obtain a significant signal to noise ratio, which increases the cost of such experiments. Further improvements can be obtained by using arbitrary waveform generator (AWG) and a higher power traveling wave tube (TWT) amplifier.

## Acknowledgements

We acknowledge the support of the Israel Science Foundation (grant 176/16 and 212/22).

**The authors declare no conflict of interest**.

