## Supporting Information for "An *in cell* site-specific labeling methodology reveals conformational changes of proteins in bacteria"

**Abstract:** Gaining new structural information on proteins in their native cellular environments will shed light on many enzymatic reaction mechanisms and encourage the development of new therapeutic approaches. During the last decade, *in cell* electron paramagnetic resonance (EPR) spectroscopy experiments have provided high-resolution data on conformational changes of proteins within the cell. However, one of the major obstacles of EPR spectroscopy is the spin-labeling process, which until now was performed only outside the cellular environment (i.e., exogenously). The spin-labeled protein is then injected into the cell, which limits the protein size and the cellular system that can be used. Here, we describe a new spin-labeling approach that can be applied to over-expressed proteins in *Escherichia coli* (i.e., endogenously). This approach uses a Cu(II) ion bound to a ligand, which has high affinity to a dHis site in the protein of interest. The presence of a nearby $^{19}$F-phenylalanine residue can be exploited to verify that the Cu(II)-ligand indeed bound to the protein target. This new methodology allows for the study of any protein, regardless of size or the cellular system used.

**Experimental Procedures**

***CueR cloning & expression protocol***

The CueR_[19]F58_L60H_G64H mutant was cloned into the pET28a expression vector by restriction-free cloning. The sequence was isolated by PCR using CueR_L60H_G64H mutant DNA as a template and the following primers: forward primer: ACC TTA CTG CGC CAG GCA CGG CAG GTG GGC TAG AAC CTG GAA GAG AGC GGC GAG CTG GTG AAT and revers primer: ATT CAC CAG CTC GCC GCT CTC TTC CAG GTT CTA GCC CAC CTG CCG TGC CTG GCG CAG TAA GGT. The clone was expressed in BL21 strain containing pDule-tfmF plasmid. A single colony containing both pET28a and pDule-tfmF vectors (ADDGENE) was used to inoculate in terrific broth (TB) medium supplemented with ampicillin and tetracycline as selection factors and grown at 37 °C to an optical density of 0.4 (at 600 nm), then 1 mM of *L-4-Trifluoromethylphenylalanine (tfmF, ALFA AESAR)* was added. The cells were grown until optical density of 0.6 and induced with 1 mM isopropyl-β-D-thiogalactopyranoside (IPTG, CALBIOCHEM) at 20°C overnight.

For in cell EPR measurements, nitrilotriacetic acid (NTA, sigma-aldrich) was incubated overnight with $CuCl_2$ at a 1:1 ratio to form a complex. The next day,150 μM of Cu(II)-NTA was added together with IPTG to *E. coli* cells that were grown until optical density of 0.6 at 600 nm. 100 mL cultures were centrifuged at 1,200 g for 20 min, washed twice with 100 mL of TB and resuspended in 200 μl of TB. 100 μL of the sample was used for in cell EPR measurements. Immediately after completing the EPR measurements, the supernatant was collected by centrifugation (2000 g at 4°C for 10 min) and its EPR spectrum was measured.

EPR measurements on lysate: The bacteria were then harvested by centrifugation at 10,000 rpm for 30 min. The pellet was re-suspended in lysis buffer (25 mM Tris, pH 7.4, 250 mM NaCl, 20 mM PMSF protease inhibitor (sigma-aldrich), 1% Triton X-100 (sigma-aldrich)) and sonicated (10 min with a duty cycle of 5 s on, 5 s off at 35% amplitude). The lysate was collected after centrifugation at 16 000 g at 4°C for 10 min.

CueR purification: The bacteria were harvested by centrifugation at 10,000 rpm for 30 min. The pellet was re-suspended in lysis buffer (25 mM Tris, pH 7.4, 250 mM NaCl, 20mM PMSF, 1% Triton X-100,) and sonicated (10 min of 30 sec pulses at 40% amplitude). The cell lysate was then centrifuged at 4°C for 30 min at 14000 rpm. Next, the protein was purified from soluble fraction by Ni-NTA agarose beads (Thermo Fisher Scientific), according to the manufacturer's protocol. The protein was eluted by addition of 250 mM imidazole to the lysis buffer. The obtained fractions were dialyzed overnight. The dialysis buffer comprised of 25 mM Tris, pH 7.4, 250 mM NaCl. The protein purity was confirmed by 12% tricine SDS-PAGE and Coomassie blue staining (Figure S1).

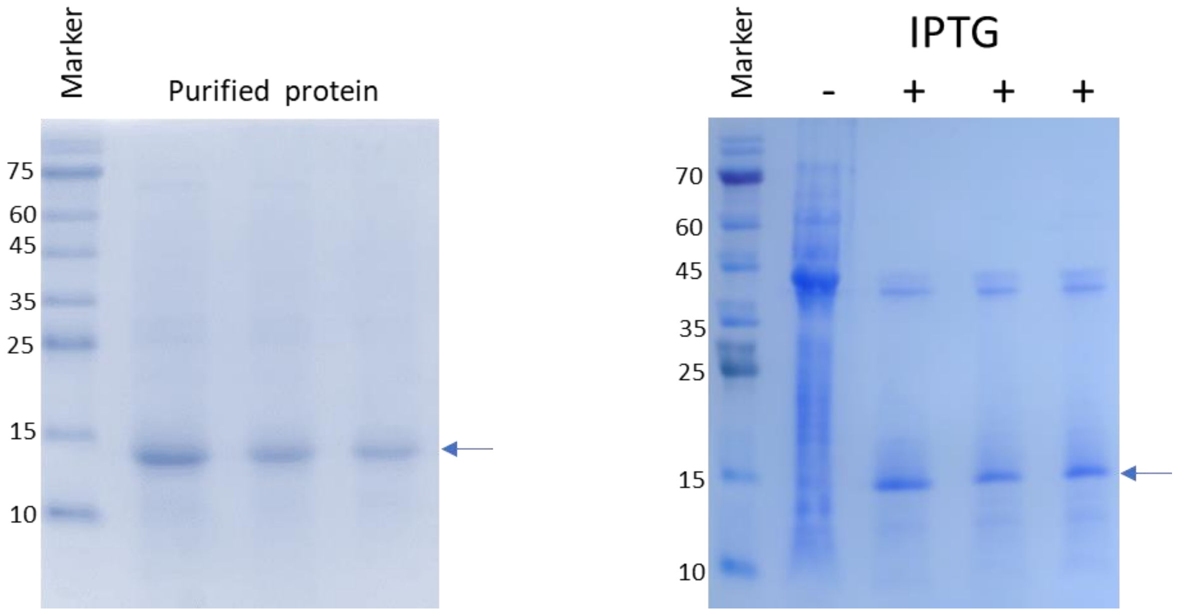


**Figure S1.** SDS-gel picture of CueR_$^{19}$F58_L60H_G64H **A**. in-cell before and after addition of IPTG and **B**. after purification. The arrow marks the 14 kDa mass of CueR.

### *Copper addition*

For in-cell EPR measurements: 50 μM $CuCl_2$ were added to the cells before measurements. For in-vitro EPR measurements on purified protein: Cu(I) (Tetrakis (acetonitrile) copper(I) hexafluorophosphate) was added to the protein solution under nitrogen gas to preserve anaerobic conditions. No Cu(II) EPR signal was observed at any time.

### *CD characterization*

Circular Dichroism (CD) measurements were performed using a Chirascan spectrometer (Applied Photophysics, UK) at room temperature. Measurements were carried out in a 1 cm optical path length cell. The data were recorded from 190–260 nm with a step size and a bandwidth of 1 nm. Spectra were obtained after background subtraction. Figure S2 shows the CD spectra of WT_CueR and CueR_$^{19}$F58_L60H_G64H mutated protein.

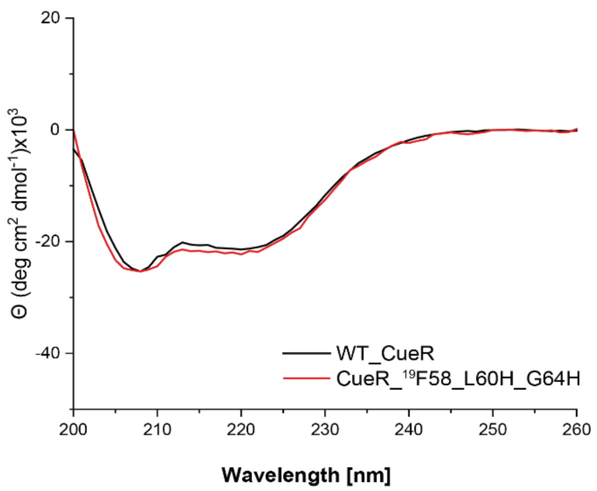


**Figure S2.** CD spectra of WT_CueR and CueR_$^{19}$F58_L60H_G64H mutated protein.

***Electrophoresis mobility shift assay (EMSA)***

EMSA experiments were carried out according to a published protocol [1]. Briefly, 5% (wt/vol) non-denaturing gels were poured and stored at 4 °C until usage. The double stranded copA promoter DNA (5'- TTGACCTTCCCCTTGCTGGAAGGTTTA -3') and protein were incubated at room temperature for 30 min. Glycerol was added to the sample reaching a final concentration of 15% (v/v) prior loading. Then, the gel was run with TAE buffer (40 mM Tris, 20 mM acetic acid, 1mM EDTA) at 4 °C, 80 V for 1 hour. Subsequently, the gel was added to an ethidium bromide staining, containing 1 µg/ml Ethidium bromide in TEA running buffer for 30 min. The gel was then washed with TEA running buffer for 30 min. The stained gel was analyzed with a Gel Doc EZ BioRad System using the EtBr protocol.

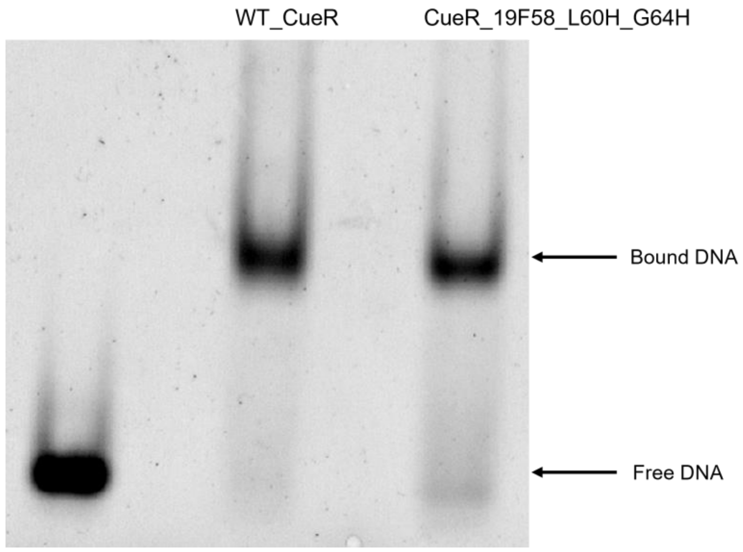


**Figure S3.** A. Electrophoresis mobility shift assay for WT_CueR and CueR_$^{19}$F58_L60H_G64H mutated protein.

**Cell Viability experiments**

BL21 strain of *E. Coli* bacteria were grown in LB at 37 °C to an optical density of 0.4 (at 600 nm), then Cu(II) or Cu(II)-NTA at various concentrations were added. Absorbance at 600 nm was measured after 16 hours.

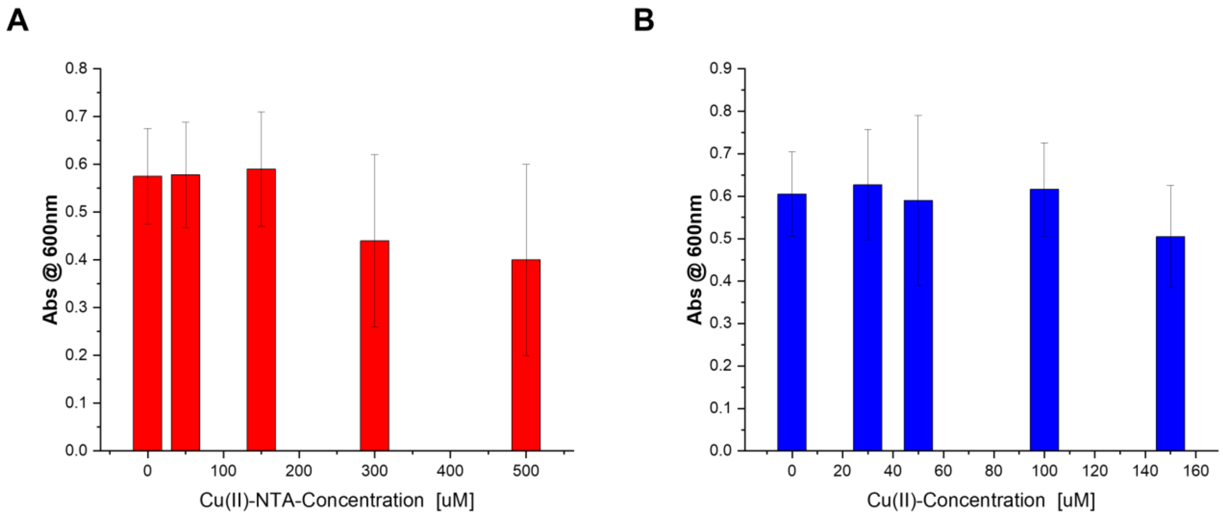


**Figure S4.** E. Coli cell viability experiments in the presence of **A**. Cu(II)-NTA and **B**. $CuCl_2$ salt.

***X-band CW EPR experiments***

Continuous-wave electron paramagnetic resonance (CW-EPR) spectra were recorded using an E500 Elexsys Bruker spectrometer operating at 9.0−9.5 GHz equipped with a super-high-sensitivity CW resonator. The spectra were recorded at low temperature (130 ± 5 K) at microwave power of 20.0 mW, modulation amplitude of 2.0 G, a time constant of 120 ms, and receiver gain of 60.0 dB. The samples were measured in 1.0-mm quartz tubes (Wilmad-LabGlass, Vineland, NJ) which was placed in 4.0 mm quartz tube for cooling process.

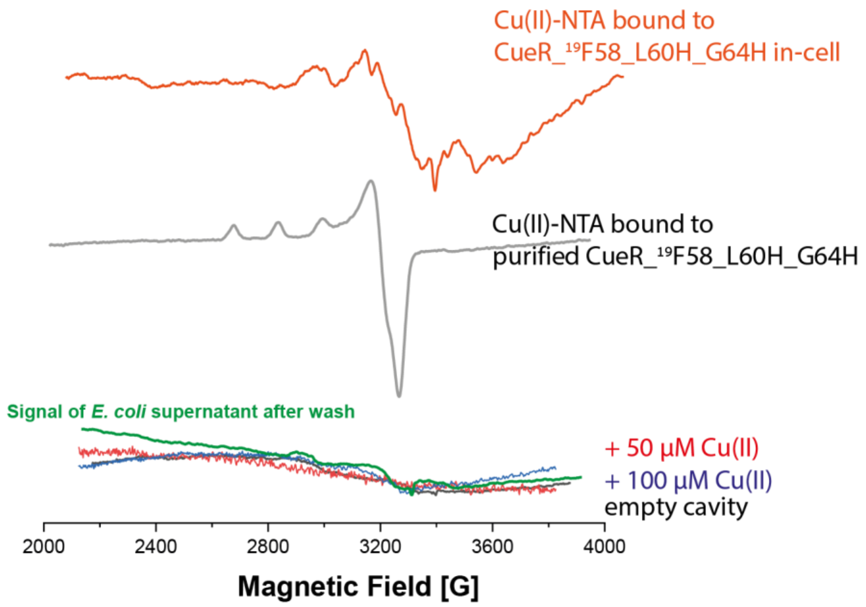


**Figure S5**. X-band CW-EPR spectra carried out at 130 ± 5 K for 50 μM (red) and 100 μM (blue) of $CuCl_2$ in *E. coli* cells, confirming that all free Cu(II) ions are reduced. The EPR spectrum of supernatant after washing the cells before EPR measurements (green). Cu(II)-NTA bound to purified CueR_$^{19}$F58_L60H_G64H (grey), and in *E. coli* cells (orange).

### *Q-band two-pulse experiments*

The two-pulse echo detected experiment were performed with π/2 pulse length of 14 ns, and a τ value of 200 ns at g⊥ position. It was carried out at 20K, 33.83GHz, 11670 G.

The two-pulse echo decay was performed with a τ value of 200 ns, and a dwell time of 16 ns.

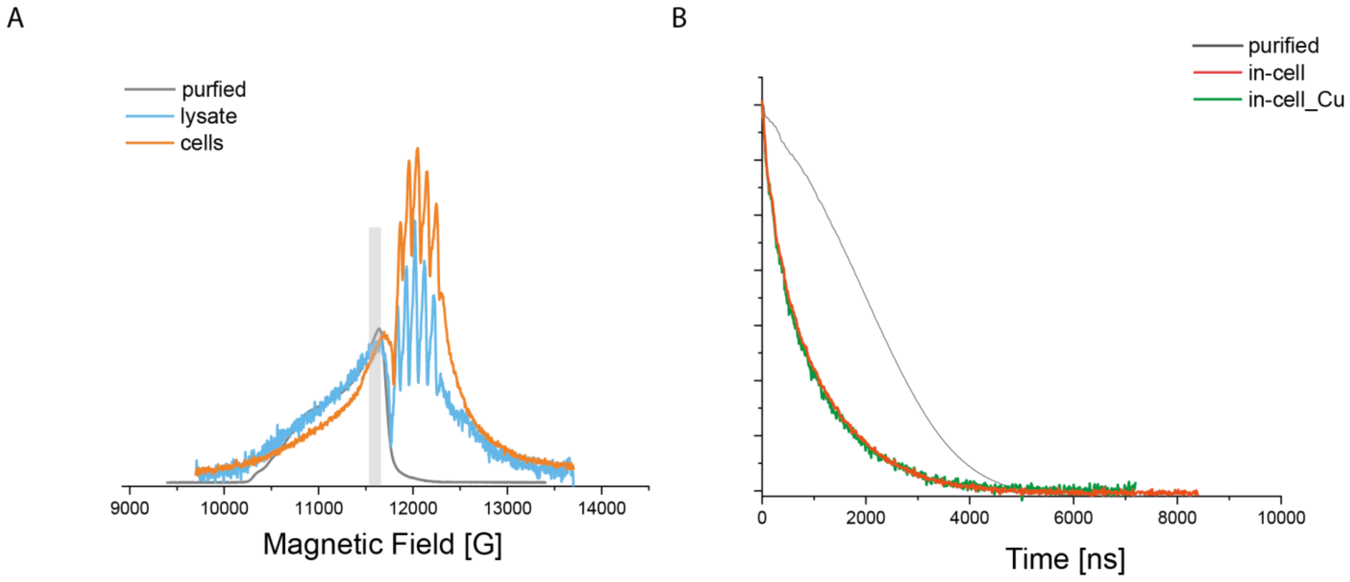


**Figure S6. Q-band two-pulse experiments A**. two-pulse field sweep of CueR_$^{19}$F58_L60H_G64H purified (grey), in lysate (light blue) and in-cell (orange). The grey rectangular marks the position where the DEER experiments were performed. **B**. two-pulse decay of CueR_$^{19}$F58_L60H_G64H purified (grey), and in-cell (with (orange) and without (green) free Cu(II) ions).

***Q-band 3P-ESEEM experiments***

The three-pulse ESEEM experiment were performed as follows: A π/2 − τ – π/2− T+dt – π/2 − τ – echo sequence was used with a four-step phase-cycle. The π/2 pulse length was 14 ns, and the τ value was set to 180 ns at $g_\perp$ position. The initial T was 300 ns, and dt was 8 ns. It was carried out at 20K, 33.83GHz, 11670 G. The data was processed by subtracting the baseline using a polynomial fit. The resulted time domain was convoluted with exponential window function, and the spectrum obtained by cross-term averaging Fourier transform after zero-fillings [2-5] (Figure S7).

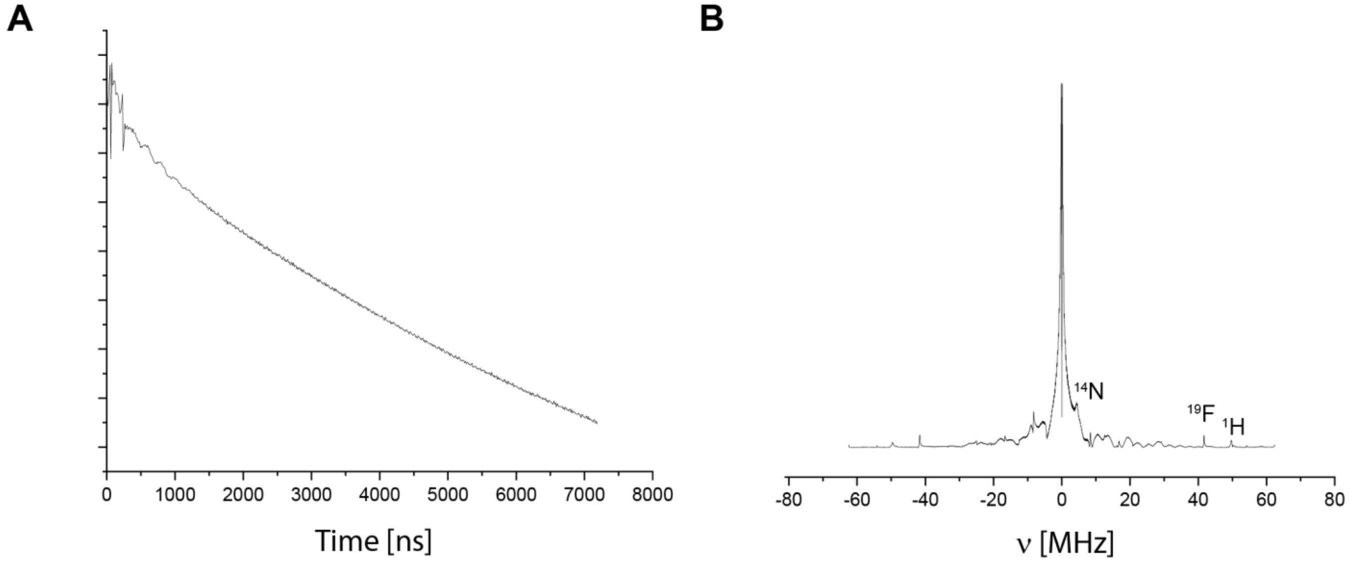


**Figure S7.** Q-band 3P-ESEEM of purified CueR_$^{19}$F58_L60H_G64H. **A**. time domain signal. **B**. FT spectrum.

***Q-band DEER measurements***

The DEER experiment $\pi/2(\nu_{obs}) - \tau_1 - \pi(\nu_{obs}) - t' - \pi(\nu_{pump}) - (\tau_1 + \tau_2 - t') - \pi(\nu_{obs}) - \tau_2 -$ echo was carried out at 50 ± 1.0 K on a Q-band Elexsys E580 spectrometer (equipped with a 2-mm probe head) and 50 W AmpQ. A two-step phase cycle was employed on the first pulse. The echo was measured as a function of t′, whereas $\tau 2$ was kept constant to eliminate relaxation effects. The durations of the observer π/2 and π pulses were 12 ns and 24 ns resecively. The duration of the π pump pulse was 24 ns, and the dwell time was 12 ns. $\tau_1$ was set to 200 ns and $\tau_2$ to 1200-1600 ns. In order to be able to compare between the various DEER signals, all data presented in the manuscript was acquired at the same frequencies and magnetic field: observer frequency: 33.83 GHz pump frequency: 33.73 GHz magnetic field: 11640 G. The samples were measured in 1.6-mm capillary quartz tubes (Wilmad-LabGlass). The DEER experiments in the cells were run for several days 3-5 days, with about 3000-5000 scans. The data was analyzed using the DeerAnalysis 2019 [6]. Both Tikhonov regularization (the regularization parameter was carefully chosen based on the best fit of the time domain and the L-curve) and DeerNet were used to analyze the data [7]. The validation considers the analysis of the two methods (Figure S9).

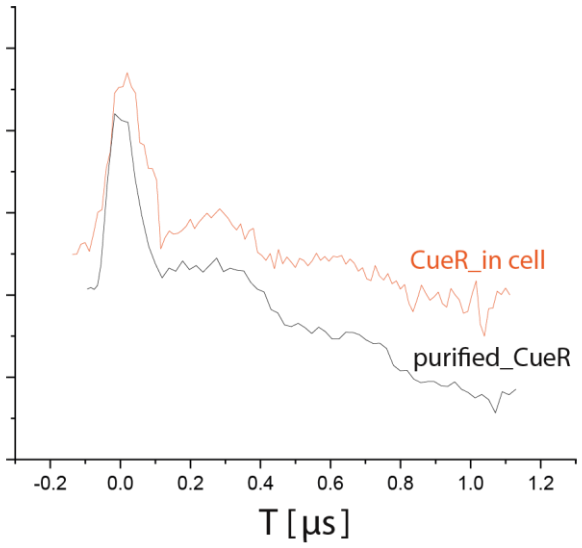


**Figure S8.** DEER time domain signals before background subtraction of purified CueR_$^{19}$F58_L60H_G64H (black) and in-cell (orange).

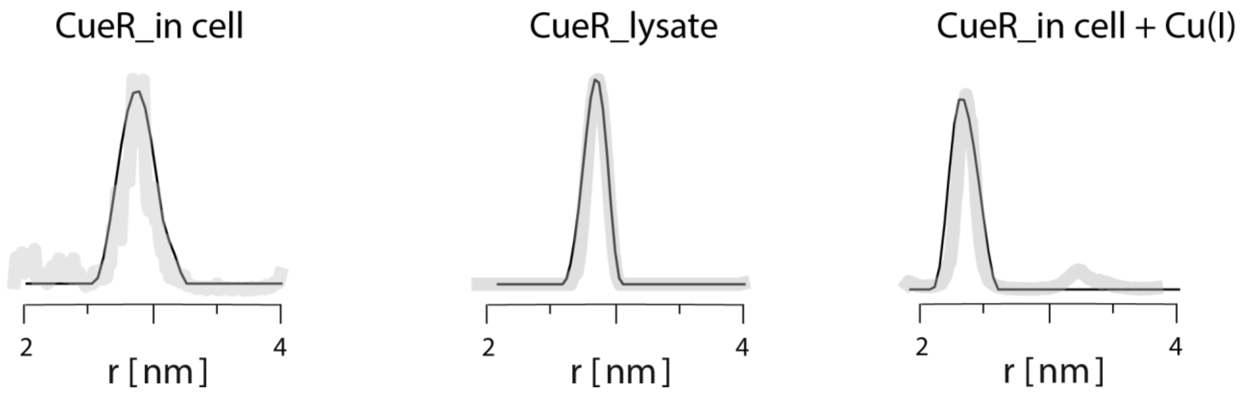


**Figure S9.** Validation Curves for the DEER distance distribution functions considering both Tikhonov regularization and DeerNET analysis.

1. Hellman, L. M., and Fried, M. G. (2007) Electrophoretic mobility shift assay (EMSA) for detecting protein-nucleic acid interactions, *Nat Protoc 2*, 1849-1861.
2. Ruthstein, S., Stone, K. M., Cunningham, T. F., Ji, M., Cascio, M., and Saxena, S. (2010) Pulsed Electron Spin Resonance Resolves the Coordination Site of Cu 2+ Ions in α1-Glycine Receptor, *Biophysical journal 99*, 2497-2506.
3. Jiang, F., McCracken, J., and Peisach, J. (1990) Nuclear quadrupole interactions in copper (II)-diethylenetriamine-substituted imidazole complexes and in copper (II) proteins, *Journal of the American Chemical Society 112*, 9035-9044.
4. Burns, C. S., Aronoff-Spencer, E., Dunham, C. M., Lario, P., Avdievich, N. I., Antholine, W. E., Olmstead, M. M., Vrielink, A., Gerfen, G. J., and Peisach, J. (2002) Molecular features of the copper binding sites in the octarepeat domain of the prion protein, *Biochemistry 41*, 3991-4001.
5. Yeagle, G. J., Gilchrist, M. L., Walker, L. M., Debus, R. J., and Britt, R. D. (2008) Multifrequency electron spin-echo envelope modulation studies of nitrogen ligation to the manganese cluster of photosystem II, *Philosophical Transactions of the Royal Society of London B: Biological Sciences 363*, 1157-1166.
6. Jeschke, G. (2007) *Deeranalysis 2006: Distance measurements on nanoscopic length scales by pulse ESR*, Springer.
7. Worswick, S. G., Spencer, J. A., Jeschke, G., and Kuprov, I. (2018) Deep neural network processing of DEER data, *Science Advances 4*.
